# Decision value signals in the ventromedial prefrontal cortex and motivational and hedonic symptoms across mood and psychotic disorders

**DOI:** 10.1101/2020.12.01.407197

**Authors:** Min Su Kang, Daniel H. Wolf, Rebecca Kazinka, Sangil Lee, Kosha Ruparel, Mark A. Elliott, Anna Xu, Matthew Cieslak, Greer Prettyman, Theodore D. Satterthwaite, Joseph W. Kable

## Abstract

Deficits in motivation and pleasure are common across many psychiatric disorders, and manifest as symptoms of amotivation and anhedonia, which are prominent features of both mood and psychotic disorders. Here we provide evidence for a shared transdiagnostic mechanism underlying impairments in motivation and pleasure across major depression, bipolar disorder, and schizophrenia. We found that value signals in the ventromedial prefrontal cortex (vmPFC) during decision-making were dampened in individuals with greater motivational and hedonic deficits, regardless of the primary diagnosis. This relationship remained significant while controlling for diagnosis-specific symptoms of mood and psychosis, such as depression as well as positive and negative symptoms. Our results demonstrate that dysfunction in the vmPFC during value-based decision-making is specifically linked to motivational and hedonic impairments across various psychiatric conditions. These findings provide a quantitative neural target for the potential development of novel treatments for amotivation and anhedonia.

---

Reductions in motivation to pursue or ability to experience pleasure, clinically termed amotivation and anhedonia, are commonly seen across a wide range of neuropsychiatric disorders and are particularly prominent in mood and psychotic disorders. These motivational and hedonic symptoms are difficult to treat and have been linked to suicidal ideation (Ballard et al., 2017; Ducasse et al., 2018), cognitive dysfunction (Franke et al., 1993; McIntyre et al., 2016), and poor clinical prognosis (McMakin et al., 2012). A growing number of theoretical papers have proposed that these symptoms may arise from a shared disruption in the brain’s reward valuation processes that occurs across many diagnostic classes of disorders (Barch et al., 2016; Husain & Roiser, 2018; Lambert et al., 2018; Whitton et al., 2015; Winograd-Gurvich et al., 2006). However, although many existing studies have compared the neural processing of rewards in cases and controls, only recently have neuroimaging studies begun to investigate motivation and pleasure using a transdiagnostic approach (Arrondo et al., 2015; Gradin et al., 2011; Park et al., 2017; Schilbach et al., 2016; Segarra et al., 2016; Sharma et al., 2017). Here we take such a transdiagnostic, dimensional framework to reveal a common circuit-level mechanism for amotivation and anhedonia across disorders, which involves disruptions in specific computational neural signals important for decision-making.

Specifically, we test the hypothesis that motivational and hedonic deficits are associated with dampened neural signals of the value of potential outcomes during decision making. This hypothesis builds on much research in decision neuroscience identifying reliable and replicable neural correlates of value during decision-making in the ventromedial prefrontal cortex (vmPFC) and ventral striatum (VS) (Bartra et al., 2013). Intriguingly, both of these regions exhibit functional and morphological abnormalities in mood and psychotic disorders (Satterthwaite et al., 2015; Wolf et al., 2014; Zhang et al., 2016). A disruption of neural value signals in these regions could lead to indifference when choosing between rewards or an unwillingness to expend effort for rewards that maps on to amotivation and anhedonia symptoms.

Our focus is on neural correlates of *decision value*, a signal about available rewards at the time a decision is made, in contrast to the focus in many previous studies on neural correlates of *experienced value*, a signal at the time a reward or conditioned cue is experienced (Platt & Plassmann, 2014). Many previous investigations of mood and psychotic disorders have measured neural activity at the time of receiving (e.g., gambling tasks (Delgado et al., 2000), guessing paradigms (Hajcak et al., 2006) or anticipating reward feedback (monetary incentive delay task (Knutson et al., 2000)). Though many of these studies have observed reduced neural signals of experienced value in response to either positive stimuli (Epstein et al., 2006; Harvey et al., 2007; Keedwell et al., 2005) or conditioned cues for positive outcomes (Juckel, Schlagenhauf, Koslowski, Filonov, et al., 2006; Juckel, Schlagenhauf, Koslowski, Wüstenberg, et al., 2006; Simon et al., 2010; Stoy et al., 2012; Wacker et al., 2009), there are also many conflicting findings (see review Zhang et al., 2016). Furthermore, accumulating evidence suggests that impairments in mood and psychotic disorders are primarily in the prospective consideration of reward, rather than in enjoyment during reward consumption (Treadway & Zald, 2011). That is, motivational and hedonic symptoms appear to primarily impact not experienced value – how people experience rewards in the moment – but rather decision value – how they evaluate potential rewards and decide which ones to pursue. Thus, considering this important distinction in decision neuroscience suggests that a disruption in decision value signals might better explain motivational and hedonic symptoms across disorders. However, to our knowledge, this hypothesis has not been investigated transdiagnostically.

To capture motivational and hedonic deficits across various classes of psychiatric disorders, we analyzed the data from 81 individuals with major depressive disorder (MDD = 17), bipolar disorder (BD = 21), schizophrenia (SCZ = 23), or no psychiatric history (HC = 20) based on the Structured Clinical Interview for DSM-IV (SCID-IV; First & Gibbon, 2004). Our primary measure of interest, symptoms of amotivation and anhedonia, was assessed using the Motivation and Pleasure (MAP) scale of the Clinical Assessment Interview for Negative Symptoms (CAINS; Kring et al., 2013). To rule out potential confounds, we also evaluated diagnosis-specific symptoms, such as depression, positive symptoms (e.g., hallucinations, delusions, disorganized speech), and negative symptoms (e.g., anhedonia, amotivation, alogia, flat affect, apathy) using the Calgary Depression Scale for Schizophrenia (CDSS; Addington et al., 1990), the Scale for the Assessment of Positive Symptoms (SAPS; Andreasen, 1989a), and the Scale for the Assessment of Negative Symptoms (SANS; Andreasen, 1989b).

We examined neural representations of decision value in these individuals in a delay discounting task, which involves choices between a smaller proximal reward and a larger reward farther in the future (Kirby & MarakoviĆ, 1996). This paradigm was one of the first used to identify neural correlates of decision value in the vmPFC and VS (Kable & Glimcher, 2007) and a recent meta-analysis confirmed that both regions reliably track decision value during the task (Schüller et al., 2019). Because the tendency to devalue future rewards varies widely across individuals, this task dissociates decision values from objective values (in this case, monetary amounts) of rewards. In addition, because this decision task specifically engages the ability to imagine the value of a future reward (i.e., no rewards are actually delivered during the task), its emphasis on the prospective evaluation of reward aligns with the clinical manifestations of amotivation and anhedonia. We expected that greater impairments in motivation and pleasure, as assessed by the CAINS MAP, would be associated with a reduced neural representation of decision values for future rewards.

## Method

### Participants

Ninety participants who met clinical eligibility were recruited from a larger study (Hershenberg et al., 2016; Wolf et al., 2014). Primary diagnosis of MDD, BD, or SCZ was ascertained using the Structured Clinical Interview for DSM-IV Axis I Disorders (SCID-IV; First & Gibbon, 2004), and all participants in the MDD and BD group were in a depressive episode at the time of the scan to minimize any variance due to manic symptoms. For BD, both type I and II were included in the study, with three participants meeting the diagnostic criteria for type II. Healthy control participants were excluded if they met criteria for any Axis I psychiatric disorder. Given the association between substance use and delay discounting, participants with a history of pathological gambling, substance abuse or dependence in the past six months (with the exception of nicotine), or a positive urine drug screen on the day of the study were excluded. Given the evidence for elevated discount rates in cigarette smokers, coupled with the higher prevalence of smoking in SCZ, all groups were matched on smoking status (Wing et al., 2012; Yu et al., 2017). Of the 90 adults who met clinical eligibility, four were excluded from the analyses due to excessive head motion during the scan, and five due to idiosyncratic responses on the task (see Quality control analysis). Therefore, our final sample consisted of 81 participants (Table 1). The excluded participants did not differ significantly on demographic and clinical variables (Supplementary Table S1). All study procedures were approved by the University of Pennsylvania’s Institutional Review Board, and all participants provided written informed consent.

**Table 1.**
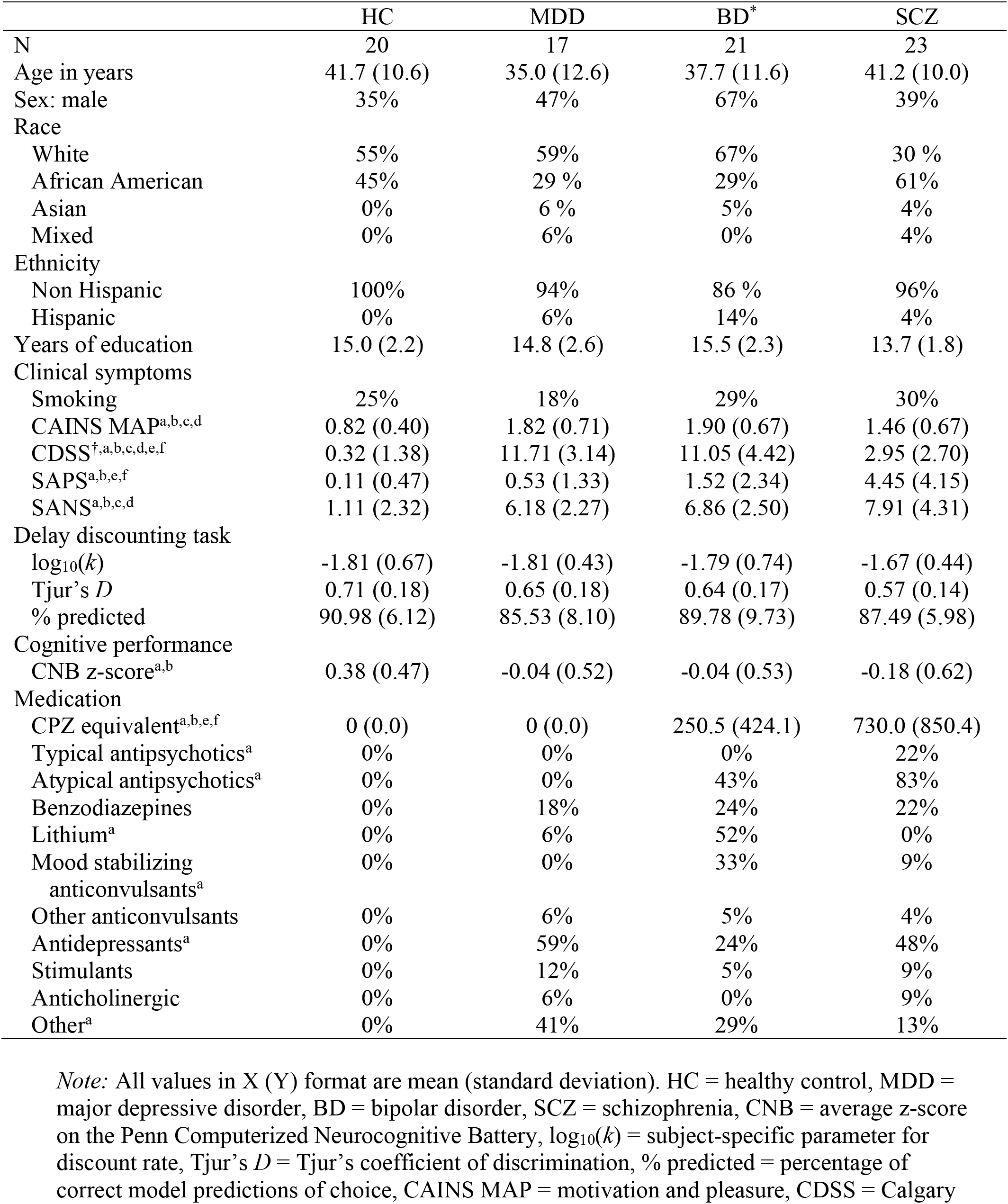
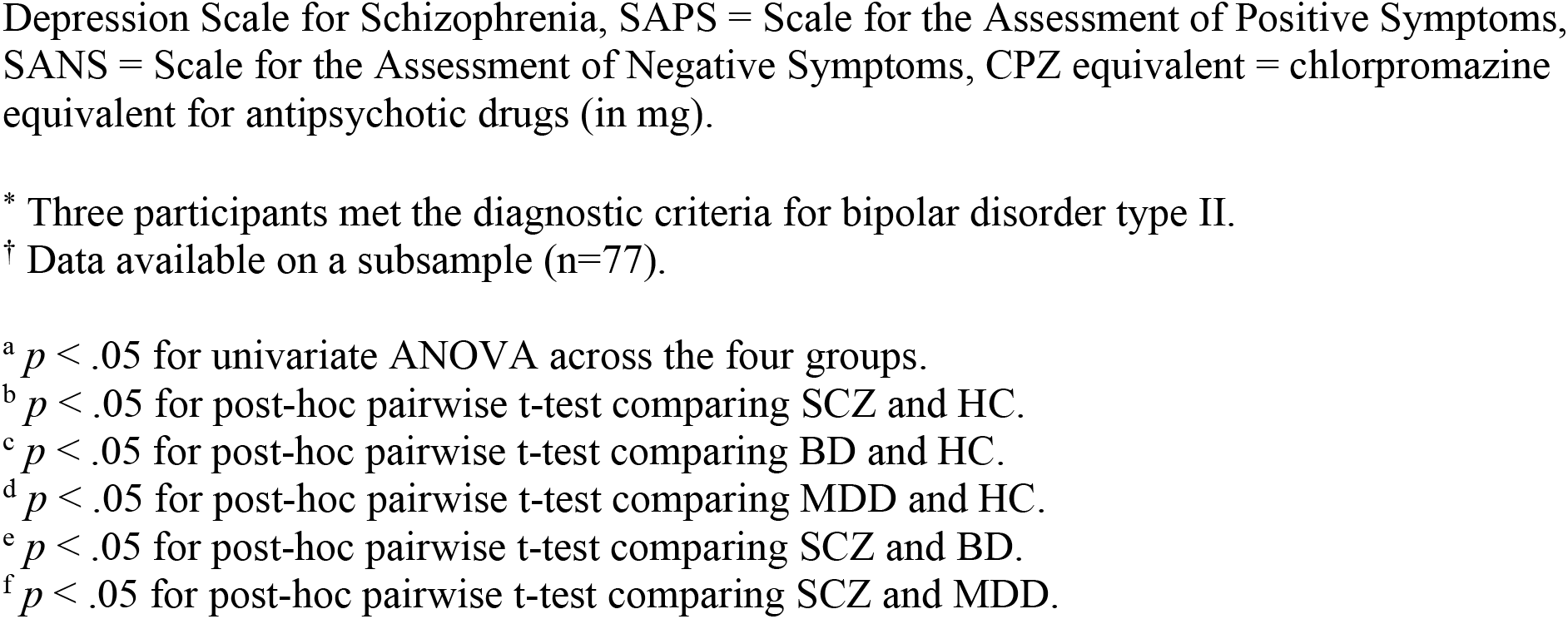
Sample characteristics.

### Clinical and cognitive measures

The Clinical Assessment Interview for Negative Symptoms - Beta (CAINS) was used as the primary measure of amotivation/anhedonia (Kring et al., 2013). The CAINS assesses motivation and pleasure in the social, recreational, and vocational domains, and therefore is less susceptible to environmental or external constraints than other existing scales (Strauss & Gold, 2016; Wolf et al., 2014). Given the two-factor structure of the CAINS (motivation and pleasure; expression), we calculated an aggregate motivation and pleasure (CAINS MAP) score by taking the mean of all relevant items. Of the 81 participants included in the analysis, 77 were also administered the Calgary Depression Scale for Schizophrenia (CDSS; Addington et al., 1990), the Scale for the Assessment of Positive Symptoms (SAPS; Andreasen, 1989a), and the Scale for the Assessment of Negative Symptoms (SANS; Andreasen, 1989b). For all clinical measures, higher scores indicate increasing levels of severity. Additionally, given the association between discount rates and cognitive abilities (Bickel et al., 2011; Heerey et al., 2011; Hinson et al., 2003), cognitive abilities were assessed using a subset of tasks from the Penn Computerized Neurocognitive Battery (CNB; Moore et al., 2015), which included (neurobehavioral function; domain): Penn Face Memory (episodic memory; face memory), Short Penn Continuous Performance Test (executive control; attention), Penn Emotion Recognition Test (social cognition; emotion identification), Penn Word Memory (episodic memory; verbal memory), Short Letter N-back (executive control; working memory), Short Penn Line Orientation Test (complex cognition; spatial ability), Short Penn Conditional Exclusion Task (executive control; mental flexibility), Short Penn Logical Reasoning Test (complex cognition, language reasoning). An aggregate cognitive functioning score was calculated by taking the mean of standardized accuracy scores (z-scores) on individual tasks.

### Delay discounting task

Participants performed a delay discounting task in the scanner (Figure 1A; Kable & Glimcher, 2010). The task consisted of 200 choices (4 runs of 50 trials) between two options: $20 now and $X in Y days, in which X ranged from $20.50 to $50 and Y ranged from 1 to 178 days. Some versions of this task vary the amount of both immediate and delayed options (Yu et al., 2017), but keeping the immediate option constant allows us to attribute variability in brain activity to changes in the decision value of the delayed reward (Kable & Glimcher, 2007, 2010). For each trial, participants had 4 s to make a response by pressing the right or left button, and the locations of the immediate and delayed options were pseudorandomized across trials. Trials in which the participant did not make a choice in 4 s were coded as missing. Each trial was followed by an inter-trial interval (ITI) that ranged from 2 to 20 s (mean ITI = 6 s). Participants were informed that one of the trials would be randomly selected for payment at the end of the experiment; payment was provided on a debit card, available immediately or after a delay according to the participant’s choice on the randomly selected trial. EPrime (https://pstnet.com) 2.0 was used for task presentation.

**Figure 1.**
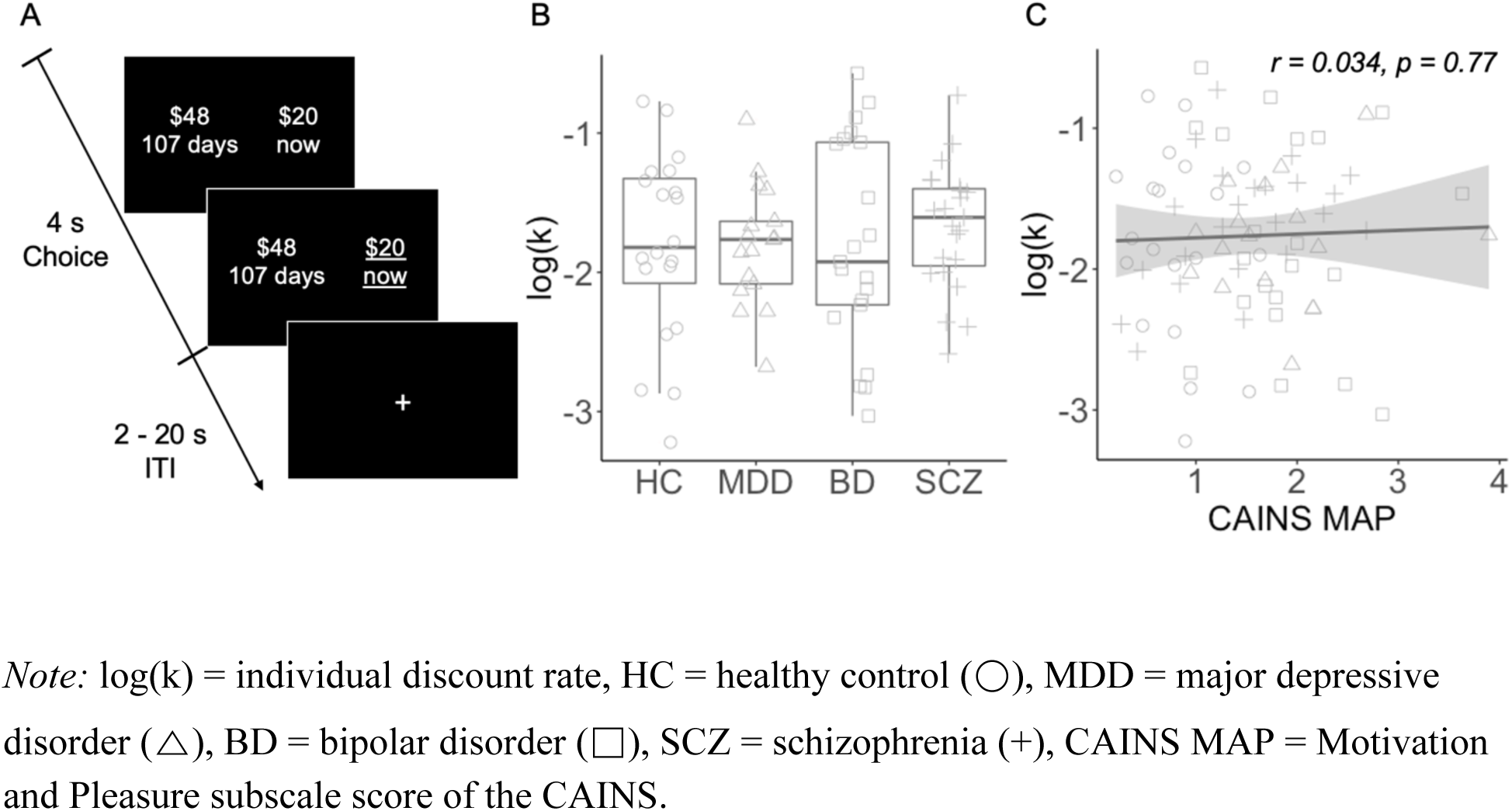
**(A)** Delay discounting fMRI paradigm. On each trial, participants chose between two options: an immediate monetary reward, fixed at $20 on every trial, and a delayed monetary reward, which varied in amount and delay across trials. Participants had 4 s to respond, and the chosen option was underlined to indicate their choice. Each trial was followed by an inter-trial interval (ITI) that ranged from 2 to 20 s. **(B)** Discount rate, log(*k*), did not differ by primary diagnosis (*F*(3,77) = 0.28, *p* = 0.84). **(C)** Discount rate was not correlated with CAINS MAP (*r* = 0.034, *p* = 0.78).

### Parameter estimation

Individual discount curves were fitted using a hyperbolic function (Kirby & Herrnstein, 1995; Lempert & Pizzagalli, 2010; Myerson & Green, 1995; Rachlin et al., 1991; Richards et al., 1999), which assumes that the decision value (DV) of the delayed reward is:

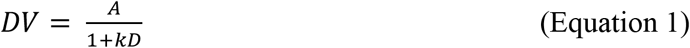

where *A* is the amount of the reward, *D* is the delay to receiving the reward, and *k* is the subject-specific free parameter for their discount rate. Participants’ individual choice data was fit with the following logistic function using maximum likelihood estimation with function minimization routines in MATLAB:

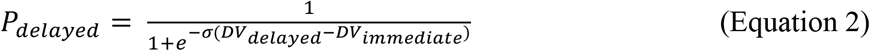

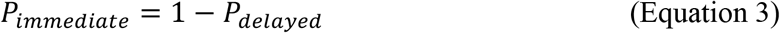

where σ is the scaling parameter in the logistic function. Because *k* was not normally distributed (Shapiro-Wilk’s *W* = 0.69, *p* < .001), individual *k* was transformed using log_10_.

### Quality control analysis

Runs that were missing more than 10% of the trials (5 out of 50 trials per run), or that were characterized by excessive head motion (see Image processing), were excluded from the analyses. Four participants were excluded because there were fewer than two runs of usable data, due to missing trials or excessive head motion. Additionally, participants were excluded if the fit of the logistic model was poor, as assessed by a Tjur’s coefficient of discrimination (Tjur’s *D*) less than 0.20. One participant was excluded due to poor model fit. Lastly, participants who chose a single option (either the immediate or delayed reward) more than 99% of the time were excluded. Three participants were excluded because they only chose the immediate option, and one participant was excluded because they only chose the delayed option. Therefore, our imaging and behavioral quality control analyses excluded 4 and 5 participants, respectively. For the subjects included for analyses (N=81), the percentage of choices of the immediate option ranged from 2.5% to 98.5% (*M* = 58.99, *SD* = 26.66).

### fMRI acquisition

All images were acquired using a Siemens Tim Trio 3T and a 32-channel head coil. For functional images, 3 mm interleaved axial slices were acquired with echo-planar T2* weighting (repetition time [TR] = 3000 ms, echo time [TE] = 30 ms, flip angle = 90°, field of view [FOV] = 192 x 192 mm, matrix = 64 x 64, slice thickness = 3 mm). Slice orientation was −30° from the anterior commissure-posterior commissure (ACPC) plane to minimize signal drop out in the orbitofrontal cortex. Each run consisted of 168 images, and the first 6 volumes (18 s) were discarded to compensate for T1 saturation effects. High-resolution T1-weighted MPRAGE anatomical images were acquired for spatial registration to a standard coordinate system (slice thickness = 1 mm, TR = 1810 ms, TE = 3.51 ms, inversion time [TI] = 1100 ms, flip angle = 9°, FOV = 1192 x 256 mm, matrix = 256 x 192, 160 slices). Additionally, for distortion correction, a B0 field map was acquired using a double-echo gradient recall echo (GRE) sequence (TR = 1000 ms, TE1 2.69 ms, TE2 5.27 ms, flip angle = 60°, FOV = 240 mm, slice thickness = 4 mm).

### Image processing

Image processing and statistical analyses were performed using FSL 5.0.9 (http://www.fmrib.ox.ac.uk/fsl). All volumes were corrected for differences in slice acquisition using Fourier-space time-series phase-shifting and corrected for small head movements using MCFLIRT (Jenkinson et al., 2002). Runs with mean relative displacement (MRD) greater than 0.30 mm were excluded. Data were smoothed using a Gaussian kernel of FWHM 6.0 mm and filtered in the temporal domain using a nonlinear high-pass filter (Gaussian-weighted least-squares straight line fitting with sigma = 50.0 s). To account for anatomical differences across subjects and to allow for statistical inference at the group level, functional images were registered to the anatomical image and spatially normalized to standard MNI space (MNI152, T1 2mm) using linear registration with FMRIB’s Linear Image Registration Tool (FLIRT) and further refined using FNIRT nonlinear registration (Andersson et al., 2007; Jenkinson et al., 2002; Jenkinson & Smith, 2001).

### fMRI analysis

Using FSL’s FMRI Expert Analysis Tool Version 6.0, we fit a general linear model (GLM) that estimated (1) averaged activity for all decisions versus rest (trial regressor) and (2) activity that was correlated across trials with the decision value of the delayed option (DV regressor, calculated using Equation 1 above with the subject-specific *k*). The first six volumes of each run were discarded prior to analysis. All other events were modeled with a fixed duration of 4 s following the stimulus presentation, and convolved with a canonical double-gamma HRF. Temporal derivatives of these two regressors, as well as the six motion parameters, were included as covariates of no interest. Missed trials, in which the participant failed to make a response in 4 s, were modeled separately.

Subsequently, all eligible runs from each participant were combined using a fixed effect model. Group-level analyses were performed using FMRIB Local Analysis of Mixed Effects module (Beckmann et al., 2003). For region-of-interest (ROI) analyses, we used vmPFC and VS masks from a meta-analysis of 206 published studies examining the neural correlates of decision value (See Figure 6A in Bartra et al., 2013). To test for any differences in activations across groups, an F-test was performed. To test for the main effect of CAINS MAP across the whole brain, individual MAP scores were demeaned and included in the GLM as an explanatory variable. Thresholded *Z* statistic images were prepared by using a threshold of *Z* > 3.1 and a corrected extent threshold of *p* < 0.05, familywise error-corrected using Gaussian Random Field Theory (Poline et al., 1997). In all multisubject statistics, outliers were de-weighted using mixture modeling (Woolrich, 2008).

### Statistical analyses

Parameter estimation and quality control analyses were performed in MATLAB R2016b (Mathworks). All imaging analyses were performed in FSL 5.0.9. All statistical analyses were performed in R 3.5.2 (CRAN). All pairwise t-tests were corrected for multiple comparisons using Holm’s method (Holm, 1979). For confound analysis, DV-related activity in the ROI was regressed on each covariate separately to identify significant covariates. All multiple linear regression models were performed to statistically control for sociodemographic variables (sex, age, education, race, and ethnicity). All categorical variables were binarized (0 or 1) and continuous variables in the regression models were z-scored to standardize effects.

## Results

### Clinical characteristics of the transdiagnostic sample

All clinical groups, regardless of the primary diagnosis, scored higher on the MAP than healthy participants (*t*_MDD>HC_ = 5.16, *t*_BD>HC_ = 6.34, *t*_SCZ>HC_ = 3.80, all *p*’s < 0.01), but did not significantly differ from one another (*F*(2,58) = 2.68, *p* = 0.08). Similarly, all clinical groups reported higher levels of overall negative symptoms than healthy participants (*t*_MDD>HC_ = 6.52, *t*_BD>HC_ = 7.44, *t*_SCZ>HC_ = 6.36, all *p*’s < 0.01), which did not differ across diagnosis (*F*(2,57) = 1.43, *p* = 0.25). Participants with mood disorders (MDD and BD) reported the highest general symptoms of depression, although those with SCZ were more depressed than healthy participants (*t*_MDD>HC_ = 13.82, *t*_BD>HC_ = 10.34, *t*_SCZ>HC_ = 4.02, all *p’s* < 0.05). Those with SCZ had more positive symptoms than the other groups (*t*_SCZ>HC_ = 4.87, *t*_SCZ>MDD_ = 4.17, *t*_SCZ>BD_ = 2.87, all *p*’s < 0.01). Additionally, SCZ also had significantly worse overall cognitive functioning than healthy participants (*t* = 3.35, *p* = 0.007). Not surprisingly, MAP was significantly correlated with total SANS negative symptoms (*r* = 0.51) and depressive symptoms (*r* = 0.54) across the entire sample. MAP was also significantly correlated with performance on the neurocognitive battery (*r* = −0.34), smoking status (*t* = 2.06) and antipsychotic medications (*r* = 0.28, all *p’s* < 0.05, Supplementary Table S2). Given these associations between MAP and other measures, we report statistical analyses evaluating these potential confounds below.

### Discount rates were not related to psychiatric diagnosis or symptoms

Discount rates, or the tendency to prefer immediate rewards, varied widely across individuals, but were not associated with primary diagnosis or MAP. Participants completed an delay discounting task that consisted of choices between $20 available now and a larger amount ($20.50-50) available in the future (1-178 days). Individual discount rates (raw *k* values) ranged from 0.0006 (preference for future rewards) to 0.27 (preference for immediate rewards), with a geometric mean of 0.017. Log-transformed discount rates (log10(*k*)) did not differ significantly across groups (*F*(3,77) = 0.28, *p* = 0.84) (Figure 1B) or by smoking status (*t* = 0.81, *p* = 0.42), and were not correlated with MAP (*r* = 0.034, *p* = 0.78) (Figure 1C). For the model estimating discount rates, we assessed two measures of model fit, Tjur’s *D* (range = 0.22−0.94, mean [SD] = 0.64 [0.17]) and the percentage of choices predicted by the model (range = 53.6%-98.7%, mean [SD] = 88.5 [7.75]). Neither measure significantly differed across groups (Tjur’s *D*: *F*(3,77) = 2.38, *p* = 0.08; % correct prediction: *F*(3,77) = 2.09, *p* = 0.11). Similarly, model fit was not significantly correlated with MAP (Tjur’s D: *r* = −0.12, *p* = 0.29; % correct prediction: *r* = −0.07, *p* = 0.54). The lack of significant differences in discount rates and model fit by diagnosis or as a function of MAP assures that any observed neural differences were not simply due to behavioral differences on the task.

### Motivational and hedonic impairments were associated with dampened value signals during decision making

Motivational and hedonic deficits were associated with dampened decision value signals in the vmPFC during the delay discounting task. Consistent with previous reports (Kable & Glimcher, 2007, 2010), decision value was correlated with activity in widespread regions, including the vmPFC, VS, and posterior cingulate cortex (PCC), across all participants (Supplementary Table S3 and Supplementary Figure S1). We defined our *a priori* regions of interest as the vmPFC and VS based on a meta-analysis of published studies examining the neural correlates of decision value (Bartra et al., 2013; Figure 2A and B). There were no significant group differences in DV-related activity in the vmPFC (*F*(3,77) = 1.84, *p* = 0.15) or VS (*F*(3,77) = 1.92, *p* = 0.13; Figure 2C and D). However, DV-related activity in the vmPFC was inversely correlated with MAP, such that individuals with increasing levels of motivational and hedonic deficits exhibited weaker value signals in the vmPFC (*r* = −0.27, *p* = 0.01; Figure 2E). The relationship between MAP and DV-related activity, though in the same direction, was not significant in the VS (*r* = −0.10, *p* = 0.36; Figure 2F).

**Figure 2.**
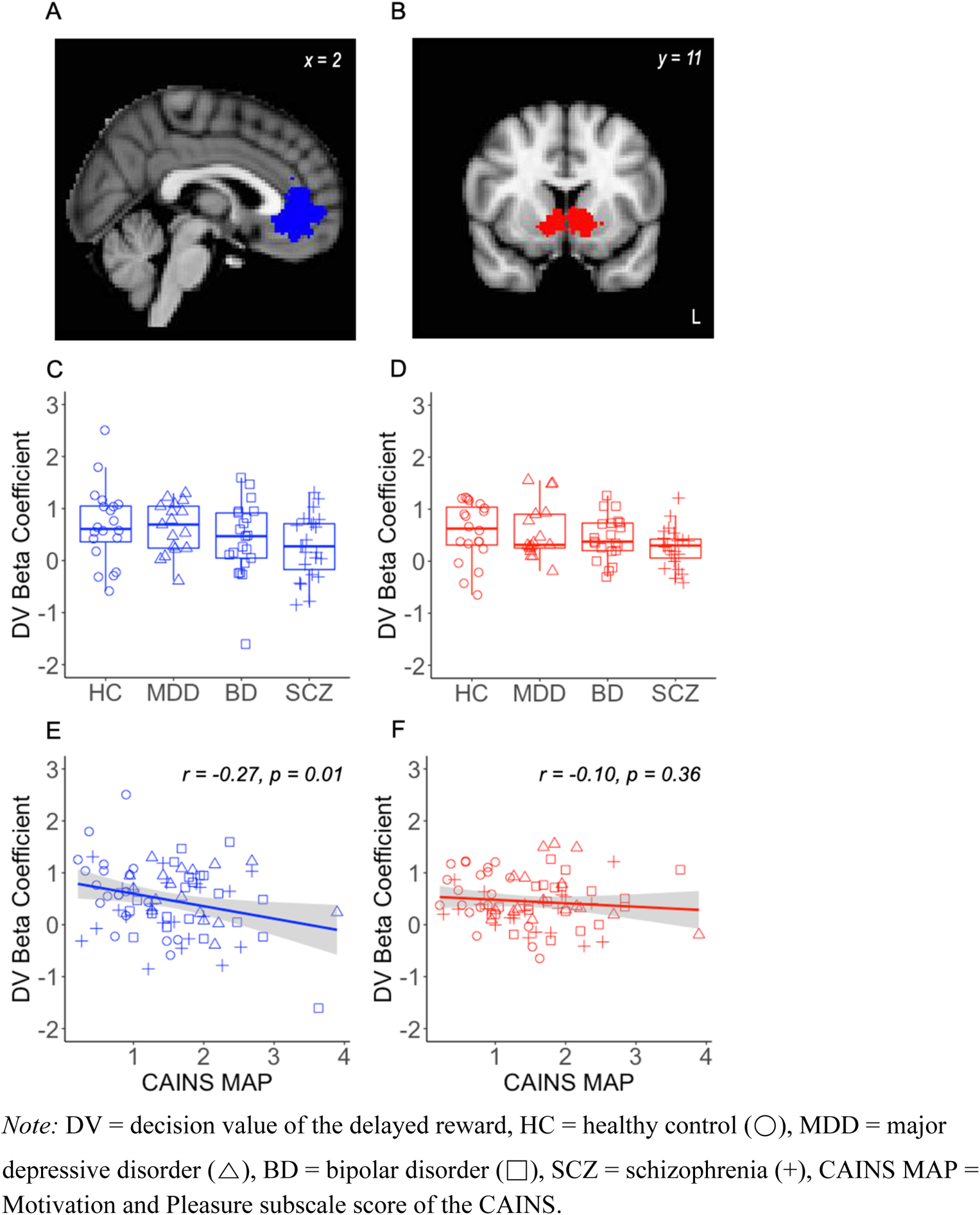
**(A, B)** Region-of-interest masks for the vmPFC (blue) and VS (red), derived from a meta-analysis by Bartra et al. (2013). There was no significant difference in DV-related activity by primary diagnosis in the **(C)** vmPFC or **(D)** VS. **(E)** DV-related activity in the vmPFC was inversely correlated with CAINS MAP. **(F)** This correlation was not significant in the VS.

### The brain-symptom relationship was specific to motivational and hedonic impairments

Though motivational and hedonic impairments were correlated with several other characteristics of the sample, these other characteristics did not explain the relationship between motivational and hedonic symptoms and reduced decision value signals in the vmPFC. To determine the specificity of the association between MAP and value-related activity in the vmPFC, we performed a series of sensitivity analyses. We first demonstrated a significant link between MAP and vmPFC activity while controlling for demographic variables such as age, sex, education, race, and ethnicity (Table 2, Model 1). We then added primary diagnosis (MDD, BD, and SCZ) to this model to rule out the possibility that the relationship is better explained by the diagnostic criteria (Table 2, Model 2). To further rule out other potential confounds, we also examined a model that includes symptoms of depression (CDSS), positive and negative symptoms (SAPS and SANS), antipsychotic medications (chlorpromazine [CPZ]-equivalent dose), performance on the neurocognitive battery (CNB), current smoking, discounting rate (log(*k*)), and model fit (Tjur’s *D*). Of these potential confounds, DV-related activity in the vmPFC was individually associated with the diagnosis of SCZ (β = −0.63, 95% CI [−1.23, - 0.03]), overall negative symptoms (β = −0.22, 95% CI [−0.44, 0.00]), CPZ-equivalent (β = −0.32, 95% CI [−0.54, −0.11]), neurocognitive performance (β = 0.27, 95% CI [0.05, 0.48]), and discounting rate (β = −0.23, 95% CI [−0.45, −0.01]). However, in multiple linear regressions that included primary diagnoses alone (Model 2) or all potential confounds (Model 3), MAP remained a significant predictor of DV-related activity, and notably, the inclusion of these covariates did not reduce the magnitude of its standardized coefficient. Because CPZ-equivalent dose primarily controls antipsychotics, we attempted to capture potential effects of other types of medications by excluding participants on any one class of medication. This series of analyses also did not significantly change the relationship between MAP and vmPFC activity (Supplementary Table S4).

**Table 2.**
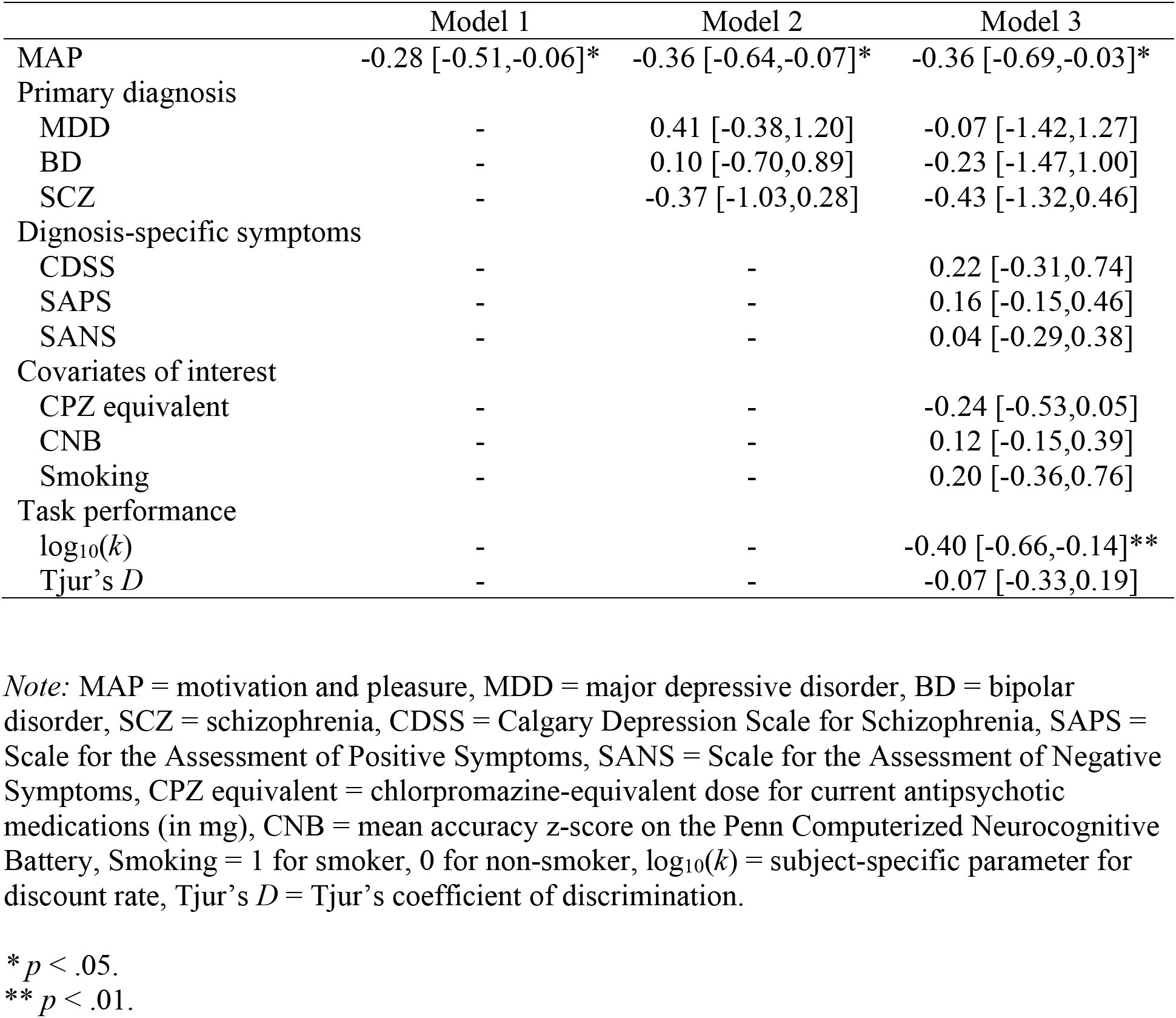
Linear regression models explaining activity in vmPFC. The dependent variable in all models was the beta coefficient for DV-related activity in the vmPFC. Beta coefficients and 95% confidence intervals for the independent variables in different models are shown. All models included sex, age, education, race, and ethnicity as covariates of no interest, and no sociodemographic covariate was significant in any model.

|  | Model 1 | Model 2 | Model 3 |
| --- | --- | --- | --- |
| MAP | -0.28 [-0.51,-0.06]* | -0.36 [-0.64,-0.07]* | -0.36 [-0.69,-0.03]* |
| Primary diagnosis |  |  |  |
| MDD | - | 0.41 [-0.38,1.20] | -0.07 [-1.42,1.27] |
| BD | - | 0.10 [-0.70,0.89] | -0.23 [-1.47,1.00] |
| SCZ | - | -0.37 [-1.03,0.28] | -0.43 [-1.32,0.46] |
| Diagnosis-specific symptoms |  |  |  |
| CDSS | - | - | 0.22 [-0.31,0.74] |
| SAPS | - | - | 0.16 [-0.15,0.46] |
| SANS | - | - | 0.04 [-0.29,0.38] |
| Covariates of interest |  |  |  |
| CPZ equivalent | - | - | -0.24 [-0.53,0.05] |
| CNB | - | - | 0.12 [-0.15,0.39] |
| Smoking | - | - | 0.20 [-0.36,0.76] |
| Task performance |  |  |  |
| $\log_{10}(k)$ | - | - | -0.40 [-0.66,-0.14]** |
| Tjur's $D$ | - | - | -0.07 [-0.33,0.19] |

*Note:* MAP = motivation and pleasure, MDD = major depressive disorder, BD = bipolar disorder, SCZ = schizophrenia, CDSS = Calgary Depression Scale for Schizophrenia, SAPS = Scale for the Assessment of Positive Symptoms, SANS = Scale for the Assessment of Negative Symptoms, CPZ equivalent = chlorpromazine-equivalent dose for current antipsychotic medications (in mg), CNB = mean accuracy z-score on the Penn Computerized Neurocognitive Battery, Smoking = 1 for smoker, 0 for non-smoker, $\log_{10}(k)$ = subject-specific parameter for discount rate, Tjur's $D$ = Tjur's coefficient of discrimination.
\* $p < .05$ .
\*\* $p < .01$ .

### Additional brain regions tracking motivational and hedonic impairments

Beyond our *a priori* ROIs, we also conducted an exploratory analysis of the correlation between MAP and decision value signals across the whole brain. We found that higher MAP scores were associated with increased DV signals in the left PCC (*Z* > 3.1, *p* < 0.05, FWE corrected; Figure 3).

**Figure 3.**
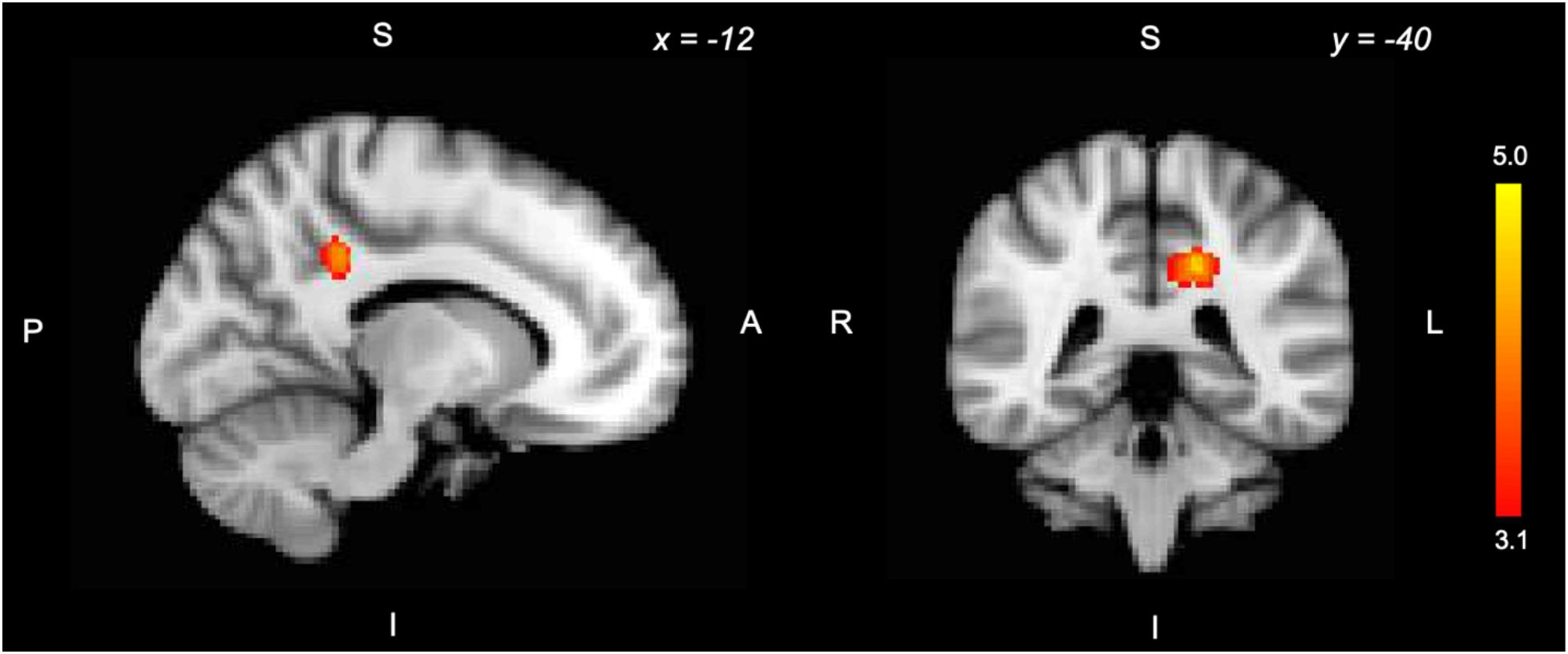
Exploratory whole-brain analysis of the correlation between DV-related activity and CAINS MAP. Higher scores on the CAINS MAP were associated with greater DV-related activity in the left PCC (*Z* > 3.1, *p* < 0.05, FWE corrected).

## Discussion

Our findings link dimensional impairments in motivation and pleasure, across mood and psychotic disorders, to disruptions in specific computational neural signals during decision-making. Dozens of previous studies have identified neural correlates of decision value, the value of potential outcomes during decision-making, in the vmPFC (see meta-analysis Bartra et al., 2013). Here we find that individuals with more severe deficits in motivation and pleasure exhibited a blunting of the decision value-related response in the vmPFC during a prospective decision-making task. Moreover, these symptoms predicted decision value signals in the vmPFC above and beyond the effects of primary diagnosis, illness-specific symptoms of depression and schizophrenia, medications, cognitive functioning, and nicotine use. These results demonstrate the specificity of the link between motivational and hedonic impairments and vmPFC function in the context of evaluating future rewards. Our findings further suggest that the vmPFC may be an important therapeutic target for amotivation and anhedonia across disorders, and demonstrate how quantitative models in decision neuroscience can help to elucidate the common pathophysiology underlying transdiagnostic, dimensional clinical deficits.

Our findings are novel and consistent with other evidence pointing to functional and morphological abnormalities in the vmPFC associated with motivational and hedonic impairments (Ward et al., 2019). Both preclinical and clinical findings suggest that vmPFC dysfunction may be normalized using direct brain stimulation, resulting in increased reward-seeking behavior in rodents and reductions of anhedonia in patients with major depression (Hamani et al., 2012; Mayberg et al., 2005). Although directly stimulating the vmPFC is not possible without surgical implants, using transcranial magnetic stimulation (TMS) or transcranial direct-current stimulation (tDCS) to indirectly stimulate the vmPFC via inter-regional functional connectivity is an emerging area of research in decision neuroscience (Figner et al., 2010; Obeso et al., 2021; Soutschek et al., 2021), and has the potential to contribute to a novel intervention that targets motivational and hedonic symptoms across disorders.

Several prior studies have examined amotivation using effort discounting, which measures the willingness to expend cognitive or physical effort for rewards (Culbreth et al., 2018). Individuals with mood and psychotic disorders tend to avoid effort expenditure (Hershenberg et al., 2016; Patzelt et al., 2019; Wolf et al., 2014), and neuroimaging studies of effort discounting have found blunted activations at the time of decision in the striatum in major depression (Yang et al., 2016) and schizophrenia (Huang et al., 20160407) and in the vmPFC in adolescents at risk for depression (Rzepa et al., 2017). An unwillingness to expend effort for rewards could be due to reduced valuation of rewards, heightened registration of effort costs, or both. An important aspect of our contribution is a test of whether amotivation is associated with dampened reward value signals during decision making, in a paradigm where there are no effort costs and amotivation is not associated with different decisions.

Several prior studies relating anhedonia to neural responses to rewards or reward cues have primarily reported blunted reward signals in the VS, and often did not find a significant effect in the vmPFC (Juckel, Schlagenhauf, Koslowski, Filonov, et al., 2006; Juckel, Schlagenhauf, Koslowski, Wüstenberg, et al., 2006; Simon et al., 2010; Stoy et al., 2012; Wacker et al., 2009). In contrast, we found that amotivation and anhedonia were associated with reduced value-related activity in the vmPFC, whereas in the VS this same relationship was a non-significant trend. This discrepancy may be partly due to the focus on *experienced* value signals in previous studies, versus the focus on *decision* value signals in the current study. Meta-analyses report stronger *experienced* value-related activity in the VS, and stronger *decision* value-related activity in the vmPFC, though both kinds of signals are present in both regions (Bartra et al., 2013; Oldham et al., 2018).

Beyond these *a priori* regions of interest, individuals with amotivation and anhedonia recruited the PCC to a greater extent in encoding the decision value of future rewards. Though exploratory, this finding is broadly consistent with a prior study that found greater PCC activation in individuals with SCZ during a delay discounting task (Avsar et al., 2013). One speculation is that this increased representation of decision value in the PCC might reflect compensation for weaker decision value signals in the vmPFC. Regardless of the interpretation, this result provides further evidence that the representation of decision values is altered in individuals with motivational and hedonic deficits.

In this study, neither diagnosis nor motivational and hedonic symptoms were associated with discount rate. Prior behavioral studies have reported elevated discount rates in SCZ (Ahn et al., 2011; Brown et al., 2018; Heerey et al., 2007, 2011; Weller et al., 2014), although results are mixed in mood disorders (Ahn et al., 2011; Brown et al., 2018; Imhoff et al., 2014; Mason et al., 2012; Pulcu et al., 2014; Takahashi et al., 2008; Urošević et al., 2016). Although not statistically significant in our sample, we observed a trend in individuals with SCZ to discount more than others. The absence of significant behavioral differences in our sample aids interpretation of the imaging findings, though, as the observed functional differences are not secondary to behavioral differences on the task. Specifically, differences in performance in clinical groups can confound the interpretation of neural differences, a limitation that is commonly addressed by matching for behavioral performance (Avsar et al., 2013; Barch et al., 2001; Cannon et al., 2005; Lee et al., 2008), although this approach often poses yet another confound as participants would be presented with different task stimuli by clinical status.

A change in discount rates, though, would not be expected to result from a disruption in decision value signals. Rather than an overall shift in preferences (in this task, a change in discount rates), dampened decision value signals would be expected to lead to an increase in the variability of choices. Lesion studies have found that vmPFC damage leads to noisier or more variable choices, rather than to overall shifts in preference (Fellows & Farah, 2007). The delay discounting task used in our study, however, was not sensitive to detecting changes in choice variability because we presented a large range of decision values to elicit robust neural effects (i.e., the majority of the decisions were easy). Therefore, we did not find any significant correlation between choice variability and MAP, although we observed a trend that all clinical groups tended to make less consistent choices compared to those without a psychiatric history. Because vmPFC signals also predict confidence during value judgments (De Martino et al., 2013), we speculate that reduced confidence in one’s decision would be another possible effect of blunted decision value signals in this region. Future studies tailoring the decision values near the participant’s indifference point (i.e., including more difficult decisions) and including trial-by-trial measures of confidence would be able to empirically test these hypotheses.

Our study has several limitations. First, the MAP subscale of the CAINS constitutes a single scale not suited for distinguishing motivation and pleasure. Anticipatory pleasure is commonly associated with motivation and goal-directed behavior, and motivation and pleasure often load on a single factor. Although psychometric, correlational, and preclinical studies suggest that clinical deficits in the two domains are interrelated, others, including the DSM, separate these into two distinct domains. We used a measure that combines motivation and pleasure because the two are clinically interrelated and our task engages both aspects of reward processing. Second, although the overall sample size was relatively large, the sample sizes for individual diagnoses were small, and therefore we did not have the sufficient statistical power to examine any within-group effect. Lastly, we did not evaluate whether vmPFC hypofunction is a general feature of decision-making, or rather specific to delay discounting. Despite these limitations, these findings demonstrate that perturbed value signals in this region during the evaluation of future rewards were specifically linked to clinical impairments in motivation and pleasure across various classes of psychiatric disorders, over and above the effects of diagnosis and general symptoms. Demonstrating such symptom-specific alterations across diagnoses helps elucidate the pathophysiological underpinnings of amotivation and anhedonia and can inform future treatment development.

## Conclusion

Taken together, our results link transdiagnostic impairments in motivation and pleasure to disruptions in specific computational neural signals during decision-making. In particular, among individuals diagnosed with mood and psychotic disorders, more severe deficits in motivation and pleasure were related to blunted decision value-related responses in the vmPFC during a prospective decision-making task. Furthermore, motivational and hedonic symptoms predicted decision value signals in this region above and beyond the effects of primary diagnosis, illness-specific symptoms of depression and schizophrenia, medications, cognitive functioning, and nicotine use. Though the causal mechanisms remain to be understood, our results suggest the therapeutic potential of interventions targeting symptom-specific changes in neural signals across transdiagnostic populations.

## Supplementary Material for

**Supplementary Table S1.** Comparison of participants included versus excluded from the analyses.

|  | Included | Excluded |
| --- | --- | --- |
| N | 81 | 9 |
| Primary diagnosis |  |  |
| HC | 20 | 3 |
| MDD | 17 | 3 |
| BD | 21 | 2 |
| SCZ | 23 | 1 |
| Sex: male | 48% | 67% |
| Race |  |  |
| White | 52% | 44% |
| African American | 42% | 56% |
| Asian | 4% | 0% |
| Mixed | 2% | 0% |
| Ethnicity |  |  |
| Non Hispanic | 94% | 100% |
| Hispanic | 6% | 0% |
| Years of education | 14.5 (2.3) | 14.6 (2.1) |
| Delay discounting task |  |  |
| $\log_{10}(k)$ | -1.76 (0.69) | -1.50 (0.73) |
| Tjur's $D$ | 0.65 (0.20) | 0.62 (0.21) |
| % predicted | 88.47 (7.75) | 86.50 (14.33) |
| Cognitive performance |  |  |
| CNB z-score | 0.02 (0.57) | 0.20 (0.64) |
| Clinical symptoms |  |  |
| Current nicotine use | 27% | 22% |
| CAINS MAP | 1.44 (0.72) | 1.64 (0.89) |
| CDSS | 5.71 (5.38) | 7.78 (8.61) |
| SAPS | 1.81 (3.10) | 1.00 (1.73) |
| SANS | 5.68 (3.96) | 5.00 (3.28) |
| Medication |  |  |
| Typical antipsychotics | 6% | 0% |
| Atypical antipsychotics | 35% | 22% |
| Benzodiazepines | 16% | 11% |
| Lithium | 15% | 11% |
| Mood stabilizing anticonvulsants | 11% | 0% |
| Other anticonvulsants | 4% | 0% |
| Antidepressants | 32% | 22% |
| Stimulants | 6% | 0% |
| Anticholinergic | 4% | 0% |
| Other | 20% | 44% |

**Supplementary Table S2.** Intercorrelation of confound variables (Pearson’s *r*).

|  | 1 | 2 | 3 | 4 | 5 | 6 | 7 |
| --- | --- | --- | --- | --- | --- | --- | --- |
| 1. CAINS MAP | - |  |  |  |  |  |  |
| 2. CNB | -0.34** | - |  |  |  |  |  |
| 3. Smoking | 0.27* | -0.20 | - |  |  |  |  |
| 4. CDSS | 0.54*** | -0.20 | 0.09 | - |  |  |  |
| 5. SAPS | 0.14 | -0.10 | 0.20 | 0.09 | - |  |  |
| 6. SANS | 0.51*** | -0.29* | 0.25* | 0.29* | 0.40*** | - |  |
| 7. CPZ-E | 0.28* | -0.21 | 0.03 | -0.09 | 0.42*** | 0.43*** | - |

*Note:* CAINS MAP = motivation and pleasure, CNB = average z-score on the Penn Computerized Neurocognitive Battery, Smoking = 1 for smoker, 0 for non-smoker, CDSS = Calgary Depression Scale for Schizophrenia, SAPS = Scale for the Assessment of Positive Symptoms, SANS = Scale for the Assessment of Negative Symptoms, CPZ-E = chlorpromazine [CPZ]-equivalent dose.
\* $p < 0.05$
\*\* $p < 0.01$
\*\*\* $p < 0.001$

**Supplementary Table S3.** Peak locations of positive effects of decision value of the delayed reward (*Z* > 3.1, *p* = 0.05, FWE corrected), corresponding to Supplemental Figure S1.

| Anatomical description | Z-value | Local maxima |  |  |
| --- | --- | --- | --- | --- |
|  |  | x | y | z |
| R. caudate | 9.32 | 6 | 12 | 2 |
| L. caudate | 9.24 | -6 | 14 | 0 |
| L. middle temporal gyrus | 8.21 | -58 | -46 | -4 |
| L. inferior frontal gyrus | 7.76 | -56 | -54 | -6 |
| L. posterior cingulate | 7.72 | 0 | -36 | 40 |
| L. ventromedial prefrontal cortex | 7.4 | -2 | 44 | 0 |

**Supplementary Table S4.**
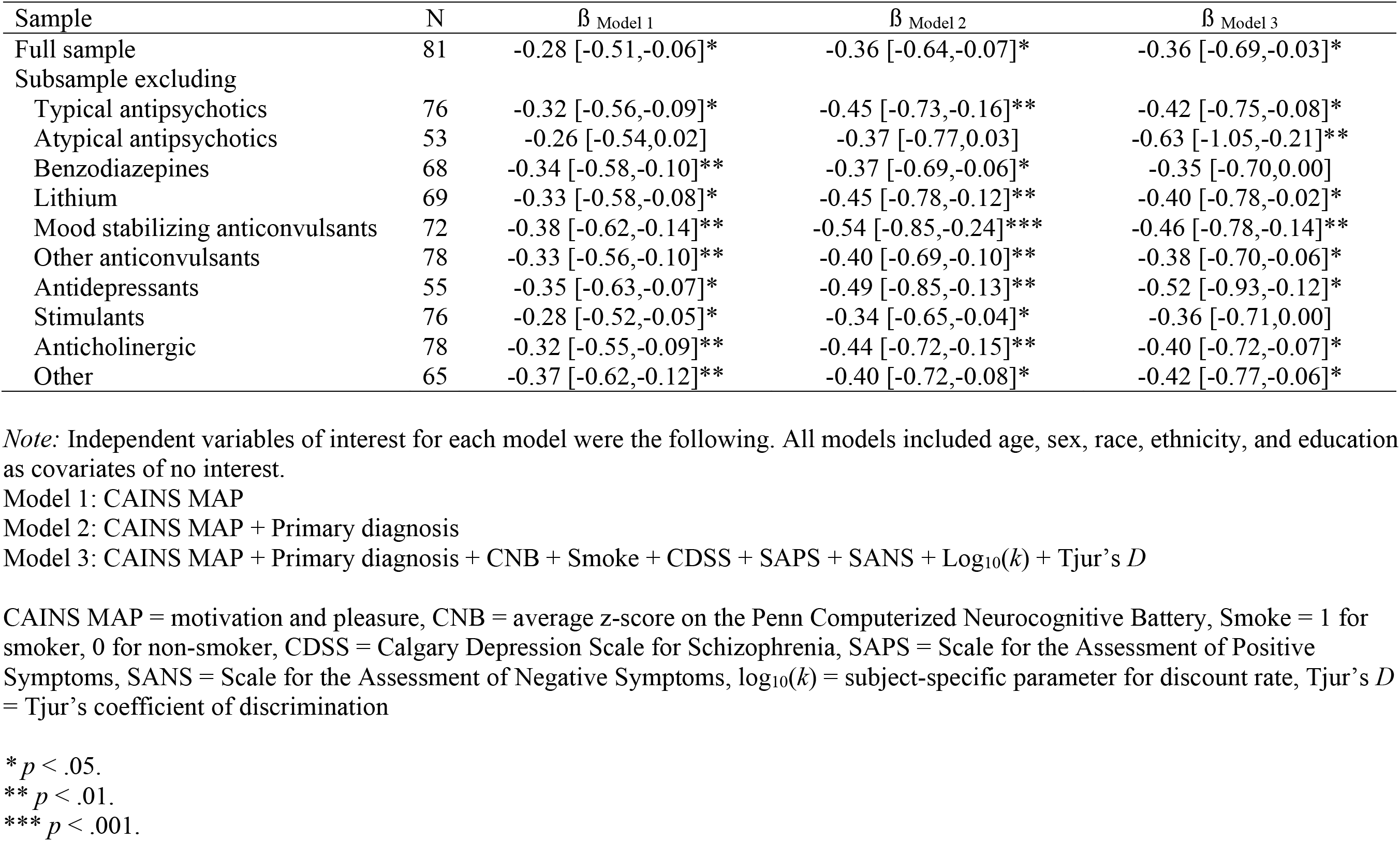
Comparison of beta coefficients and standard errors for CAINS MAP from linear regression models in subsamples excluding participants on each class of medications (dependent variable = beta coefficients for DV in the vmPFC).

**Supplemental Figure S1.**
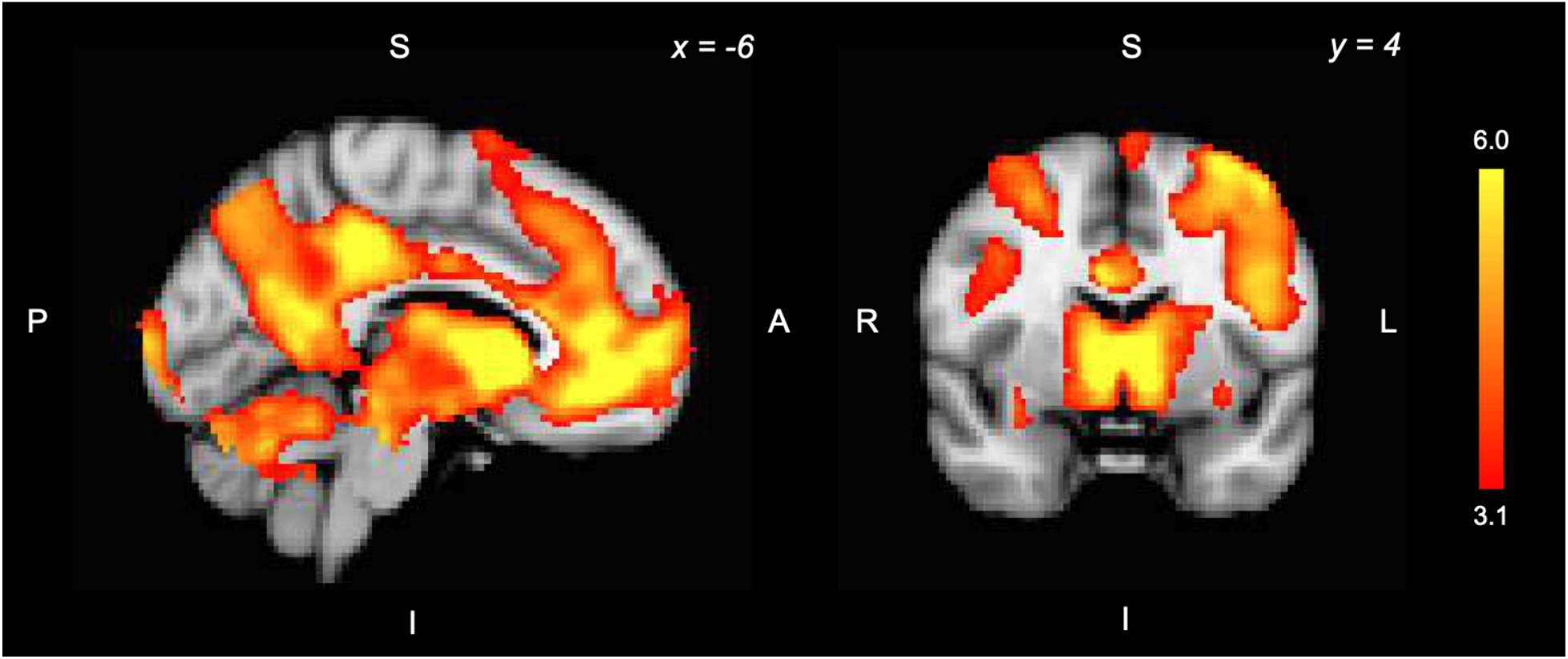
Positive effect of decision value. Across all participants, the decision value of delayed rewards was correlated with activity in widespread regions, including vmPFC, VS, and PCC (*Z* > 3.1, *p* = 0.05, FWE corrected).

## References

Addington, D., Addington, J., & Schissel, B. (1990). A depression rating scale for schizophrenics. Schizophrenia Research, 3(4), 247–251.

Ahn, W.-Y., Rass, O., Fridberg, D. J., Bishara, A. J., Forsyth, J. K., Breier, A., Busemeyer, J. R., Hetrick, W. P., Bolbecker, A. R., & O’Donnell, B. F. (2011). Temporal discounting of rewards in patients with bipolar disorder and schizophrenia. Journal of Abnormal Psychology, 120(4), 911–921. https://doi.org/10.1037/a0023333

Andersson, J. L., Jenkinson, M., & Smith, S. (2007). Non-linear registration, aka spatial normalisation FMRIB Technial Report TR07JA2. FMRIB Analysis Group of the University of Oxford.

Andreasen, N. C. (1989a). Scale for the Assessment of Positive Symptoms (SAPS). The British Journal of Psychiatry, 155(Suppl 7), 53–58.

Andreasen, N. C. (1989b). The Scale for the Assessment of Negative Symptoms (SANS): Conceptual and Theoretical Foundations. The British Journal of Psychiatry, 155(S7), 49–52. https://doi.org/10.1192/S0007125000291496

Arrondo, G., Segarra, N., Metastasio, A., Ziauddeen, H., Spencer, J., Reinders, N. R., Dudas, R. B., Robbins, T. W., Fletcher, P. C., & Murray, G. K. (2015). Reduction in ventral striatal activity when anticipating a reward in depression and schizophrenia: A replicated cross-diagnostic finding. Frontiers in Psychology, 6. https://doi.org/10.3389/fpsyg.2015.01280

Avsar, K. B., Weller, R. E., Cox, J. E., Reid, M. A., White, D. M., & Lahti, A. C. (2013). An fMRI investigation of delay discounting in patients with schizophrenia. Brain and Behavior, 3(4), 384–401. https://doi.org/10.1002/brb3.135

Ballard, E. D., Wills, K., Lally, N., Richards, E. M., Luckenbaugh, D. A., Walls, T., Ameli, R., Niciu, M. J., Brutsche, N. E., Park, L., & Zarate, J. C. (2017). Anhedonia as a clinical correlate of suicidal thoughts in clinical ketamine trials. Journal of Affective Disorders, 218, 195–200. https://doi.org/10.1016/j.jad.2017.04.057

Barch, D. M., Carter, C. S., Braver, T. S., Sabb, F. W., MacDonald, A., Noll, D. C., & Cohen, J. D. (2001). Selective Deficits in Prefrontal Cortex Function in Medication-Naive Patients With Schizophrenia. Archives of General Psychiatry, 58(3), 280–288. https://doi.org/10.1001/archpsyc.58.3.280

Barch, D. M., Pagliaccio, D., & Luking, K. (2016). Mechanisms Underlying Motivational Deficits in Psychopathology: Similarities and Differences in Depression and Schizophrenia. In E. H. Simpson & P. D. Balsam (Eds.), Behavioral Neuroscience of Motivation (pp. 411–449). Springer International Publishing. https://doi.org/10.1007/7854_2015_376

Bartra, O., McGuire, J. T., & Kable, J. W. (2013). The valuation system: A coordinate-based meta-analysis of BOLD fMRI experiments examining neural correlates of subjective value. NeuroImage, 76, 412–427. https://doi.org/10.1016/j.neuroimage.2013.02.063

Beckmann, C. F., Jenkinson, M., & Smith, S. M. (2003). General multilevel linear modeling for group analysis in FMRI. NeuroImage, 20(2), 1052–1063. https://doi.org/10.1016/S1053-8119(03)00435-X

Bickel, W. K., Yi, R., Landes, R. D., Hill, P. F., & Baxter, C. (2011). Remember the Future: Working Memory Training Decreases Delay Discounting Among Stimulant Addicts. Biological Psychiatry, 69(3), 260–265. https://doi.org/10.1016/j.biopsych.2010.08.017

Brown, H. E., Hart, K. L., Snapper, L. A., Roffman, J. L., & Perlis, R. H. (2018). Impairment in delay discounting in schizophrenia and schizoaffective disorder but not primary mood disorders. Npj Schizophrenia, 4(1), 9. https://doi.org/10.1038/s41537-018-0050-z

Cannon, T. D., Glahn, D. C., Kim, J., Erp, T. G. M. V., Karlsgodt, K., Cohen, M. S., Nuechterlein, K. H., Bava, S., & Shirinyan, D. (2005). Dorsolateral Prefrontal Cortex Activity During Maintenance and Manipulation of Information in Working Memory in Patients With Schizophrenia. Archives of General Psychiatry, 62(10), 1071–1080. https://doi.org/10.1001/archpsyc.62.10.1071

Culbreth, A. J., Moran, E. K., & Barch, D. M. (2018). Effort-cost decision-making in psychosis and depression: Could a similar behavioral deficit arise from disparate psychological and neural mechanisms? Psychological Medicine, 48(6), 889–904. https://doi.org/10.1017/S0033291717002525

De Martino, B., Fleming, S. M., Garrett, N., & Dolan, R. (2013). Confidence in value-based choice. Nature Neuroscience, 16(1), 105–110. https://doi.org/10.1038/nn.3279

Delgado, M. R., Nystrom, L. E., Fissell, C., Noll, D. C., & Fiez, J. A. (2000). Tracking the hemodynamic responses to reward and punishment in the striatum. Journal of Neurophysiology, 84(6), 3072–3077. https://doi.org/10.1152/jn.2000.84.6.3072

Ducasse, D., Loas, G., Dassa, D., Gramaglia, C., Zeppegno, P., Guillaume, S., Olié, E., & Courtet, P. (2018). Anhedonia is associated with suicidal ideation independently of depression: A meta-analysis. Depression and Anxiety, 35(5), 382–392. https://doi.org/10.1002/da.22709

Epstein, J., Pan, H., Kocsis, J. H., Yang, Y., Butler, T., Chusid, J., Hochberg, H., Murrough, J., Strohmayer, E., Stern, E., & Silbersweig, D. A. (2006). Lack of ventral striatal response to positive stimuli in depressed versus normal subjects. The American Journal of Psychiatry, 163(10), 1784–1790. https://doi.org/10.1176/ajp.2006.163.10.1784

Fellows, L. K., & Farah, M. J. (2007). The Role of Ventromedial Prefrontal Cortex in Decision Making: Judgment under Uncertainty or Judgment Per Se? Cerebral Cortex, 17(11), 2669–2674. https://doi.org/10.1093/cercor/bhl176

Figner, B., Knoch, D., Johnson, E. J., Krosch, A. R., Lisanby, S. H., Fehr, E., & Weber, E. U. (2010). Lateral prefrontal cortex and self-control in intertemporal choice. Nature Neuroscience, 13(5), 538–539. https://doi.org/10.1038/nn.2516

First, M. B., & Gibbon, M. (2004). The Structured Clinical Interview for DSM-IV Axis I Disorders (SCID-I) and the Structured Clinical Interview for DSM-IV Axis II Disorders (SCID-II). In Comprehensive handbook of psychological assessment, Vol. 2: Personality assessment (pp. 134–143). John Wiley & Sons Inc.

Franke, P., Maier, W., Hardt, J., & Hain, C. (1993). Cognitive functioning and anhedonia in subjects at risk for schizophrenia. Schizophrenia Research, 10(1), 77–84. https://doi.org/10.1016/0920-9964(93)90079-X

Gradin, V. B., Kumar, P., Waiter, G., Ahearn, T., Stickle, C., Milders, M., Reid, I., Hall, J., & Steele, J. D. (2011). Expected value and prediction error abnormalities in depression and schizophrenia. Brain: A Journal of Neurology, 134(Pt 6), 1751–1764. https://doi.org/10.1093/brain/awr059

Hajcak, G., Moser, J. S., Holroyd, C. B., & Simons, R. F. (2006). The feedback-related negativity reflects the binary evaluation of good versus bad outcomes. Biological Psychology, 71(2), 148–154. https://doi.org/10.1016/j.biopsycho.2005.04.001

Hamani, C., Machado, D. C., Hipólide, D. C., Dubiela, F. P., Suchecki, D., Macedo, C. E., Tescarollo, F., Martins, U., Covolan, L., & Nobrega, J. N. (2012). Deep Brain Stimulation Reverses Anhedonic-Like Behavior in a Chronic Model of Depression: Role of Serotonin and Brain Derived Neurotrophic Factor. Biological Psychiatry, 71(1), 30–35. https://doi.org/10.1016/j.biopsych.2011.08.025

Harvey, P.-O., Pruessner, J., Czechowska, Y., & Lepage, M. (2007). Individual differences in trait anhedonia: A structural and functional magnetic resonance imaging study in non-clinical subjects. Molecular Psychiatry, 12(8), 703, 767–775. https://doi.org/10.1038/sj.mp.4002021

Heerey, E. A., Matveeva, T. M., & Gold, J. M. (2011). Imagining the future: Degraded representations of future rewards and events in schizophrenia. Journal of Abnormal Psychology, 120(2), 483–489. https://doi.org/10.1037/a0021810

Heerey, E. A., Robinson, B. M., McMahon, R. P., & Gold, J. M. (2007). Delay discounting in schizophrenia. Cognitive Neuropsychiatry, 12(3), 213–221. https://doi.org/10.1080/13546800601005900

Hershenberg, R., Satterthwaite, T. D., Daldal, A., Katchmar, N., Moore, T. M., Kable, J. W., & Wolf, D. H. (2016). Diminished effort on a progressive ratio task in both unipolar and bipolar depression. Journal of Affective Disorders, 196, 97–100. https://doi.org/10.1016/j.jad.2016.02.003

Hinson, J. M., Jameson, T. L., & Whitney, P. (2003). Impulsive decision making and working memory. Journal of Experimental Psychology. Learning, Memory, and Cognition, 29(2), 298–306.

Holm, S. (1979). A Simple Sequentially Rejective Multiple Test Procedure. Scandinavian Journal of Statistics, 6(2), 65–70. JSTOR.

Huang, J., Yang, X., Lan, Y., Zhu, C., Liu, X., Wang, Y., Cheung, E. F. C., Xie, G., & Chan, R. C. K. (20160407). Neural substrates of the impaired effort expenditure decision making in schizophrenia. Neuropsychology, 30(6), 685. https://doi.org/10.1037/neu0000284

Husain, M., & Roiser, J. P. (2018). Neuroscience of apathy and anhedonia: A transdiagnostic approach. Nature Reviews Neuroscience, 19(8), 470–484. https://doi.org/10.1038/s41583-018-0029-9

Imhoff, S., Harris, M., Weiser, J., & Reynolds, B. (2014). Delay discounting by depressed and non-depressed adolescent smokers and non-smokers. Drug and Alcohol Dependence, 135, 152–155. https://doi.org/10.1016/j.drugalcdep.2013.11.014

Jenkinson, M., Bannister, P., Brady, M., & Smith, S. (2002). Improved Optimization for the Robust and Accurate Linear Registration and Motion Correction of Brain Images. NeuroImage, 17(2), 825–841. https://doi.org/10.1006/nimg.2002.1132

Jenkinson, M., & Smith, S. (2001). A global optimisation method for robust affine registration of brain images. Medical Image Analysis, 5(2), 143–156.

Juckel, G., Schlagenhauf, F., Koslowski, M., Filonov, D., Wüstenberg, T., Villringer, A., Knutson, B., Kienast, T., Gallinat, J., Wrase, J., & Heinz, A. (2006). Dysfunction of ventral striatal reward prediction in schizophrenic patients treated with typical, not atypical, neuroleptics. Psychopharmacology, 187(2), 222–228. https://doi.org/10.1007/s00213-006-0405-4

Juckel, G., Schlagenhauf, F., Koslowski, M., Wüstenberg, T., Villringer, A., Knutson, B., Wrase, J., & Heinz, A. (2006). Dysfunction of ventral striatal reward prediction in schizophrenia. NeuroImage, 29(2), 409–416. https://doi.org/10.1016/j.neuroimage.2005.07.051

Kable, J. W., & Glimcher, P. W. (2007). The neural correlates of subjective value during intertemporal choice. Nature Neuroscience, 10(12), 1625–1633. https://doi.org/10.1038/nn2007

Kable, J. W., & Glimcher, P. W. (2010). An “as soon as possible” effect in human intertemporal decision making: Behavioral evidence and neural mechanisms. Journal of Neurophysiology, 103(5), 2513–2531. https://doi.org/10.1152/jn.00177.2009

Keedwell, P. A., Andrew, C., Williams, S. C. R., Brammer, M. J., & Phillips, M. L. (2005). The neural correlates of anhedonia in major depressive disorder. Biological Psychiatry, 58(11), 843–853. https://doi.org/10.1016/j.biopsych.2005.05.019

Kirby, K. N., & Herrnstein, R. J. (1995). Preference Reversals Due to Myopic Discounting of Delayed Reward. Psychological Science, 6(2), 83–89. https://doi.org/10.1111/j.1467-9280.1995.tb00311.x

Kirby, K. N., & MarakoviĆ, N. N. (1996). Delay-discounting probabilistic rewards: Rates decrease as amounts increase. Psychonomic Bulletin & Review, 3(1), 100–104. https://doi.org/10.3758/BF03210748

Knutson, B., Westdorp, A., Kaiser, E., & Hommer, D. (2000). FMRI Visualization of Brain Activity during a Monetary Incentive Delay Task. NeuroImage, 12(1), 20–27. https://doi.org/10.1006/nimg.2000.0593

Kring, A. M., Gur, R. E., Blanchard, J. J., Horan, W. P., & Reise, S. P. (2013). The Clinical Assessment Interview for Negative Symptoms (CAINS): Final Development and Validation. The American Journal of Psychiatry, 170(2), 165–172. https://doi.org/10.1176/appi.ajp.2012.12010109

Lambert, C., Da Silva, S., Ceniti, A. K., Rizvi, S. J., Foussias, G., & Kennedy, S. H. (2018). Anhedonia in depression and schizophrenia: A transdiagnostic challenge. CNS Neurosci Ther., 24, 615–623. https://doi.org/10.1111/cns.12854

Lee, J., Folley, B. S., Gore, J., & Park, S. (2008). Origins of Spatial Working Memory Deficits in Schizophrenia: An Event-Related fMRI and Near-Infrared Spectroscopy Study. PLOS ONE, 3(3), e1760. https://doi.org/10.1371/journal.pone.0001760

Lempert, K. M., & Pizzagalli, D. A. (2010). Delay Discounting and Future-directed Thinking in Anhedonic Individuals. Journal of Behavior Therapy and Experimental Psychiatry, 41(3), 258–264. https://doi.org/10.1016/j.jbtep.2010.02.003

Mason, L., O’Sullivan, N., Blackburn, M., Bentall, R., & El-Deredy, W. (2012). I want it now! Neural correlates of hypersensitivity to immediate reward in hypomania. Biological Psychiatry, 71(6), 530–537. https://doi.org/10.1016/j.biopsych.2011.10.008

Mayberg, H. S., Lozano, A. M., Voon, V., McNeely, H. E., Seminowicz, D., Hamani, C., Schwalb, J. M., & Kennedy, S. H. (2005). Deep Brain Stimulation for Treatment-Resistant Depression. Neuron, 45(5), 651–660. https://doi.org/10.1016/j.neuron.2005.02.014

McIntyre, R. S., Woldeyohannes, H. O., Soczynska, J. K., Maruschak, N. A., Wium-Andersen, I. K., Vinberg, M., Cha, D. S., Lee, Y., Xiao, H. X., Gallaugher, L. A., Dale, R. M., Alsuwaidan, M. T., Mansur, R. B., Muzina, D. J., Carvalho, A. F., Jerrell, J. M., & Kennedy, S. H. (2016). Anhedonia and cognitive function in adults with MDD: Results from the International Mood Disorders Collaborative Project. CNS Spectrums, 21(5), 362–366. https://doi.org/10.1017/S1092852915000747

McMakin, D. L., Olino, T. M., Porta, G., Dietz, L. J., Emslie, G., Clarke, G., Wagner, K. D., Asarnow, J. R., Ryan, N. D., Birmaher, B., Shamseddeen, W., Mayes, T., Kennard, B., Spirito, A., Keller, M., Lynch, F. L., Dickerson, J. F., & Brent, D. A. (2012). Anhedonia Predicts Poorer Recovery Among Youth With Selective Serotonin Reuptake Inhibitor Treatment–Resistant Depression. Journal of the American Academy of Child & Adolescent Psychiatry, 51(4), 404–411. https://doi.org/10.1016/j.jaac.2012.01.011

Moore, T. M., Reise, S. P., Gur, R. E., Hakonarson, H., & Gur, R. C. (2015). Psychometric properties of the Penn Computerized Neurocognitive Battery. Neuropsychology, 29(2), 235–246. https://doi.org/10.1037/neu0000093

Myerson, J., & Green, L. (1995). Discounting of Delayed Rewards: Models of Individual Choice. Journal of the Experimental Analysis of Behavior, 64(3), 263–276. https://doi.org/10.1901/jeab.1995.64-263

Obeso, I., Herrero, M.-T., Ligneul, R., Rothwell, J. C., & Jahanshahi, M. (2021). A Causal Role for the Right Dorsolateral Prefrontal Cortex in Avoidance of Risky Choices and Making Advantageous Selections. Neuroscience, 458, 166–179. https://doi.org/10.1016/j.neuroscience.2020.12.035

Oldham, S., Murawski, C., Fornito, A., Youssef, G., Yücel, M., & Lorenzetti, V. (2018). The anticipation and outcome phases of reward and loss processing: A neuroimaging meta-analysis of the monetary incentive delay task. Human Brain Mapping, 39(8), 3398–3418. https://doi.org/10.1002/hbm.24184

Park, I. H., Lee, B. C., Kim, J.-J., Kim, J. I., & Koo, M.-S. (2017). Effort-Based Reinforcement Processing and Functional Connectivity Underlying Amotivation in Medicated Patients with Depression and Schizophrenia. Journal of Neuroscience, 37(16), 4370–4380. https://doi.org/10.1523/JNEUROSCI.2524-16.2017

Patzelt, E. H., Kool, W., Millner, A. J., & Gershman, S. J. (2019). The transdiagnostic structure of mental effort avoidance. Scientific Reports, 9(1), 1689. https://doi.org/10.1038/s41598-018-37802-1

Platt, M. L., & Plassmann, H. (2014). Chapter 13—Multistage Valuation Signals and Common Neural Currencies. In P. W. Glimcher & E. Fehr (Eds.), Neuroeconomics (Second Edition) (pp. 237–258). Academic Press. https://doi.org/10.1016/B978-0-12-416008-8.00013-9

Poline, J. B., Worsley, K. J., Evans, A. C., & Friston, K. J. (1997). Combining spatial extent and peak intensity to test for activations in functional imaging. NeuroImage, 5(2), 83–96. https://doi.org/10.1006/nimg.1996.0248

Pulcu, E., Trotter, P. D., Thomas, E. J., McFarquhar, M., Juhasz, G., Sahakian, B. J., Deakin, J. F. W., Zahn, R., Anderson, I. M., & Elliott, R. (2014). Temporal discounting in major depressive disorder. Psychological Medicine, 44(9), 1825–1834. https://doi.org/10.1017/S0033291713002584

Rachlin, H., Raineri, A., & Cross, D. (1991). Subjective probability and delay. Journal of the Experimental Analysis of Behavior, 55(2), 233–244. https://doi.org/10.1901/jeab.1991.55-233

Richards, J. B., Zhang, L., Mitchell, S. H., & Wit, H. de. (1999). Delay or Probability Discounting in a Model of Impulsive Behavior: Effect of Alcohol. Journal of the Experimental Analysis of Behavior, 71(2), 121–143. https://doi.org/10.1901/jeab.1999.71-121

Rzepa, E., Fisk, J., & McCabe, C. (2017). Blunted neural response to anticipation, effort and consummation of reward and aversion in adolescents with depression symptomatology. Journal of Psychopharmacology (Oxford, England), 31(3), 303–311. https://doi.org/10.1177/0269881116681416

Satterthwaite, T. D., Kable, J. W., Vandekar, L., Katchmar, N., Bassett, D. S., Baldassano, C. F., Ruparel, K., Elliott, M. A., Sheline, Y. I., Gur, R. C., Gur, R. E., Davatzikos, C., Leibenluft, E., Thase, M. E., & Wolf, D. H. (2015). Common and Dissociable Dysfunction of the Reward System in Bipolar and Unipolar Depression. Neuropsychopharmacology: Official Publication of the American College of Neuropsychopharmacology, 40(9), 2258–2268. https://doi.org/10.1038/npp.2015.75

Schilbach, L., Hoffstaedter, F., Müller, V., Cieslik, E. C., Goya-Maldonado, R., Trost, S., Sorg, C., Riedl, V., Jardri, R., Sommer, I., Kogler, L., Derntl, B., Gruber, O., & Eickhoff, S. B. (2016). Transdiagnostic commonalities and differences in resting state functional connectivity of the default mode network in schizophrenia and major depression. NeuroImage: Clinical, 10, 326–335. https://doi.org/10.1016/j.nicl.2015.11.021

Schüller, C. B., Kuhn, J., Jessen, F., & Hu, X. (2019). Neuronal correlates of delay discounting in healthy subjects and its implication for addiction: An ALE meta-analysis study. The American Journal of Drug and Alcohol Abuse, 45(1), 51–66. https://doi.org/10.1080/00952990.2018.1557675

Segarra, N., Metastasio, A., Ziauddeen, H., Spencer, J., Reinders, N. R., Dudas, R. B., Arrondo, G., Robbins, T. W., Clark, L., Fletcher, P. C., & Murray, G. K. (2016). Abnormal Frontostriatal Activity During Unexpected Reward Receipt in Depression and Schizophrenia: Relationship to Anhedonia. Neuropsychopharmacology, 41(8), 2001–2010. https://doi.org/10.1038/npp.2015.370

Sharma, A., Wolf, D. H., Ciric, R., Kable, J. W., Moore, T. M., Vandekar, S. N., Katchmar, N., Daldal, A., Ruparel, K., Davatzikos, C., Elliott, M. A., Calkins, M. E., Shinohara, R. T., Bassett, D. S., & Satterthwaite, T. D. (2017). Connectome-Wide Analysis Reveals Common Dimensional Reward Deficits Across Mood and Psychotic Disorders. The American Journal of Psychiatry, 174(7), 657–666. https://doi.org/10.1176/appi.ajp.2016.16070774

Simon, J. J., Biller, A., Walther, S., Roesch-Ely, D., Stippich, C., Weisbrod, M., & Kaiser, S. (2010). Neural correlates of reward processing in schizophrenia—Relationship to apathy and depression. Schizophrenia Research, 118(1), 154–161. https://doi.org/10.1016/j.schres.2009.11.007

Soutschek, A., Moisa, M., Ruff, C. C., & Tobler, P. N. (2021). Frontopolar theta oscillations link metacognition with prospective decision making. Nature Communications, 12(1), 3943. https://doi.org/10.1038/s41467-021-24197-3

Stoy, M., Schlagenhauf, F., Sterzer, P., Bermpohl, F., Hägele, C., Suchotzki, K., Schmack, K., Wrase, J., Ricken, R., Knutson, B., Adli, M., Bauer, M., Heinz, A., & Ströhle, A. (2012). Hyporeactivity of ventral striatum towards incentive stimuli in unmedicated depressed patients normalizes after treatment with escitalopram. Journal of Psychopharmacology, 26(5), 677–688. https://doi.org/10.1177/0269881111416686

Strauss, G. P., & Gold, J. M. (2016). A Psychometric Comparison of the Clinical Assessment Interview for Negative Symptoms and the Brief Negative Symptom Scale. Schizophrenia Bulletin, 42(6), 1384–1394. https://doi.org/10.1093/schbul/sbw046

Takahashi, T., Oono, H., Inoue, T., Boku, S., Kako, Y., Kitaichi, Y., Kusumi, I., Masui, T., Nakagawa, S., Suzuki, K., Tanaka, T., Koyama, T., & Radford, M. H. B. (2008). Depressive patients are more impulsive and inconsistent in intertemporal choice behavior for monetary gain and loss than healthy subjects—An analysis based on Tsallis’ statistics. Neuro Endocrinology Letters, 29(3), 351–358.

Treadway, M. T., & Zald, D. H. (2011). Reconsidering anhedonia in depression: Lessons from translational neuroscience. Neuroscience & Biobehavioral Reviews, 35(3), 537–555. https://doi.org/10.1016/j.neubiorev.2010.06.006

Urošević, S., Youngstrom, E. A., Collins, P., Jensen, J. B., & Luciana, M. (2016). Associations of Age with Reward Delay Discounting and Response Inhibition in Adolescents with Bipolar Disorders. Journal of Affective Disorders, 190, 649–656. https://doi.org/10.1016/j.jad.2015.11.005

Wacker, J., Dillon, D. G., & Pizzagalli, D. A. (2009). The role of the nucleus accumbens and rostral anterior cingulate cortex in anhedonia: Integration of resting EEG, fMRI, and volumetric techniques. NeuroImage, 46(1), 327–337. https://doi.org/10.1016/j.neuroimage.2009.01.058

Ward, J., Lyall, L. M., Bethlehem, R. A. I., Ferguson, A., Strawbridge, R. J., Lyall, D. M., Cullen, B., Graham, N., Johnston, K. J. A., Bailey, M. E. S., Murray, G. K., & Smith, D. J. (2019). Novel genome-wide associations for anhedonia, genetic correlation with psychiatric disorders, and polygenic association with brain structure. Translational Psychiatry, 9(1), 327. https://doi.org/10.1038/s41398-019-0635-y

Weller, R. E., Avsar, K. B., Cox, J. E., Reid, M. A., White, D. M., & Lahti, A. C. (2014). Delay discounting and task performance consistency in patients with schizophrenia. Psychiatry Research, 215(2), 286–293. https://doi.org/10.1016/j.psychres.2013.11.013

Whitton, A., Treadway, M., & Pizzagalli, D. (2015). Reward processing dysfunction in major depression, bipolar disorder and schizophrenia. Current Opinion in Psychiatry, 28(1), 7–12. https://doi.org/10.1097/YCO.0000000000000122

Wing, V. C., Moss, T. G., Rabin, R. A., & George, T. P. (2012). Effects of cigarette smoking status on delay discounting in schizophrenia and healthy controls. Addictive Behaviors, 37(1), 67–72. https://doi.org/10.1016/j.addbeh.2011.08.012

Winograd-Gurvich, C., Fitzgerald, P. B., Georgiou-Karistianis, N., Bradshaw, J. L., & White, O. B. (2006). Negative symptoms: A review of schizophrenia, melancholic depression and Parkinson’s disease. Brain Research Bulletin, 70(4), 312–321. https://doi.org/10.1016/j.brainresbull.2006.06.007

Wolf, D. H., Satterthwaite, T. D., Kantrowitz, J. J., Katchmar, N., Vandekar, L., Elliott, M. A., & Ruparel, K. (2014). Amotivation in Schizophrenia: Integrated Assessment With Behavioral, Clinical, and Imaging Measures. Schizophrenia Bulletin, 40(6), 1328–1337. https://doi.org/10.1093/schbul/sbu026

Woolrich, M. (2008). Robust group analysis using outlier inference. NeuroImage, 41(2), 286–301. https://doi.org/10.1016/j.neuroimage.2008.02.042

Yang, X., Huang, J., Lan, Y., Zhu, C., Liu, X., Wang, Y., Cheung, E. F. C., Xie, G., & Chan, R. C. K. (2016). Diminished caudate and superior temporal gyrus responses to effort-based decision making in patients with first-episode major depressive disorder. Progress in Neuro-Psychopharmacology and Biological Psychiatry, 64, 52–59. https://doi.org/10.1016/j.pnpbp.2015.07.006

Yu, L. Q., Lee, S., Katchmar, N., Satterthwaite, T. D., Kable, J. W., & Wolf, D. H. (2017). Steeper discounting of delayed rewards in schizophrenia but not first-degree Relatives. Psychiatry Research, 252, 303–309. https://doi.org/10.1016/j.psychres.2017.02.062

Zhang, B., Lin, P., Shi, H., Öngür, D., Auerbach, R. P., Wang, X., Yao, S., & Wang, X. (2016). Mapping anhedonia-specific dysfunction in a transdiagnostic approach: An ALE meta-analysis. Brain Imaging and Behavior, 10(3), 920–939. https://doi.org/10.1007/s11682-015-9457-6

